# Discovery of a stress-response integrative and conjugative element from *Sphingopyxis granuli* TFA broadly conserved across Sphingomonadales and Rhizobiales

**DOI:** 10.64898/2025.12.11.693706

**Authors:** Inmaculada García-Romero, Gabriel Santos-Díaz, Antonio Moreno-Rodríguez, Juan A. Marqués-Hinojosa, Antonio J. Pérez-Pulido, Francisca Reyes-Ramírez, Alejandro Rubio

## Abstract

Horizontal gene transfer is a pivotal element in the evolution of microbes, enabling them to acquire novel genes and phenotypes. Integrative and conjugative elements (ICEs) are a type of mobile genetic element that can integrate into the host genome and propagate during chromosome replication and cell division. The induction of ICE gene expression results in the excision of the ICE gene, the production of conserved conjugation machinery, a Type IV Secretion System, and the potential for DNA transfer to appropriate receptors. It has been observed that ICEs frequently contain cargo genes that do not typically align with the ICE life cycle. These genes often result in the manifestation of phenotypes that are of particular interest. The bacterium *Sphingopyxis granuli* strain TFA is being studied for its ability to degrade the contaminant tetralin present in crude oils. Genomic analysis identified eight possible integrative mobile elements in *S. granuli* TFA. Most of these regions exhibited a distribution pattern that was restricted to the species, and they lacked some functional modules that are characteristic of a complete ICE. This finding suggests the presence of degenerate structures or limited mobilization capacity. However, only two of the detected elements exhibited the capacity to retain all the modules necessary for transfer, integration, and maintenance. These elements also contained a cargo module that included genes associated with lipid metabolic pathways and resistance mechanisms. Among them, ICE3 was distinguished as the sole complete functional ICE that was also present in other species. Transcriptomic analysis under multiple stress conditions revealed differential and consistent activation of ICE3 genes, demonstrating their direct contribution to bacterial resilience and suggesting a key adaptive role in response to adverse environmental changes.

## Introduction

Bacteria possess a remarkable capacity for evolution and adaptation, as evidenced by their ability to acquire genes through horizontal transfer. This process contributes to the diversification of their genomes and facilitates the development of novel phenotypes (Tokuda & Shintani, 2024). There are three well-studied mechanisms of horizontal gene transfer: transformation, the natural ability of bacteria to absorb exogenous DNA from the environment; transduction, the transfer of DNA from one cell to another by means of a bacteriophage; and conjugation, the unidirectional, cell-to-cell contact-dependent transfer of DNA from a donor to a recipient via a conjugation apparatus expressed in the donor. Both transduction and conjugation involve mobile genetic elements, which can mediate their own transfer from one cell to another. Other types of mobile genetic elements, such as transposons and insertion sequences, are mobile within an organism, but not necessarily between organisms (Johnson & Grossman, 2015).

The primary transfer vehicles comprise plasmids, phages, transposons, and integrative and conjugative elements (ICEs). These represent a particularly sophisticated form of genetic exchange, combining the stability of chromosomally integrated elements with the dissemination capacity of conjugative plasmids (Wozniak & Waldor, 2010). These modular entities possess the capacity to integrate stably into the bacterial chromosome, replicate along with it, and, under certain physiological or environmental signals, excise and transfer to another host via a type IV conjugation system (Haskett et al., 2017).

ICEs are organized into conserved functional modules that encode proteins of interest for their function. The absence of any of these modules may prevent the naming and detection of the ICE (Bean et al., 2022; Delavat et al., 2017). Among these characteristics, the following stand out: Firstly, DNA regions are identified that are flanked by recombination sites, designated as attL and attR. Secondly, these regions undergo a process of recombination within the chromosome, facilitated by the activity of an integrase, a site-specific recombinase. This process enables the excision and subsequent integration of DNA from the donor bacterium into the host chromosome. Thirdly, a relaxase, termed Mob, is present. This relaxase is a transesterase enzyme that functions in the initiation of the process of mobilization and conjugation by nicking the origin of transfer (oriT). Finally, a series of genes encoding a type IV secretion system (T4SS) is observed, which facilitates horizontal transfer (Cury et al., 2017). This transfer module has been linked to conjugative plasmid transfer systems, and to the pathogenicity secretion systems of *Agrobacterium tumefaciens* or *Helicobacter pylori* (Christie et al., 2014). Consequently, these mobile elements exhibit characteristics of conjugative plasmids, including the type IV secretion system (T4SS) and the T4CP coupling protein, while also employing integration mechanisms characteristic of prophages, facilitated by tyrosine- or serine-type integrases or recombinases.

In addition to these essential components, it should be noted that ICEs incorporate highly variable accessory regions where genes that confer specific adaptive advantages to their hosts reside (Gomberg & Grossman, 2024). Numerous studies have demonstrated the pivotal role of ICEs in ecological adaptation and bacterial metabolic innovation. In clinical contexts, these elements have been significant contributors to the spread of antibiotic resistance or virulence factors. This phenomenon has been reported in hypervirulent and multidrug-resistant strains of *Klebsiella pneumoniae* that cause infections in humans and *Pasteurella multocida*, which also affects farm animals (He et al., 2024; Russo & Marr, 2019). Notably, the ICE SXT/R391 of *Vibrio cholerae* is key to the global spread of antimicrobial resistance genes (Wozniak et al., 2009). In symbiotic bacteria, such as *Mesorhizobium loti* and *Bradyrhizobium japonicum*, ICEs carrying nitrogen fixation and nodulation genes have been described, showing how these elements promote mutualistic plant-bacteria associations (Sullivan & Ronson, 1998). Finally, there are examples of ECEs involved in the degradation of aromatic pollutants, such as ICEclc from *Pseudomonas knackmussii* B13, which contains genes involved in the degradation of chlorinated aromatic compounds. This contributes to the adaptation of this bacterium to contaminated soils and plays a key role in bioremediation (Sentchilo et al., 2009). These examples reflect the evolutionary plasticity of ICEs and their role as dynamic platforms for genetic exchange.

The order Sphingomonadales, which includes genera such as *Sphingomonas*, *Sphingobium*, *Novosphingobium*, and *Sphingopyxis*, represents a group of Gram-negative bacteria with a remarkable ability to degrade aromatic compounds derived from petroleum. These compounds include benzene, naphthalene, tetralin, and phenanthrene (Asaf et al., 2020). These bacteria are frequently found in contaminated soils, wastewater, and industrial ecosystems, where they actively contribute to the biotransformation of organic pollutants. It has been demonstrated that various species of the genera *Sphingomonas*, *Sphingopyxis*, and *Sphingobium* possess highly dynamic genomes that are abundant in mobile elements. Consequently, the metabolic plasticity exhibited by Sphingomonadales has been linked to the prevalence of mobile genetic elements, including plasmids, genomic islands, and ICEs. These elements facilitate the transfer and reorganization of catabolic operons and resistance systems (Song et al., 2023). In this context, *Sphingopyxis granuli* TFA is a noteworthy exemplar of metabolic adaptation. This particular strain, isolated from a hydrocarbon-enriched bioreactor, has demonstrated a remarkable ability to utilize tetralin as its exclusive source of carbon and energy. Tetralin, an aromatic compound present in petroleum fractions and industrial pollutants, serves as the strain’s primary carbon and energy source (García-Romero et al., 2016, 2020). Its metabolic versatility includes the ability to grow under microaerobic conditions and tolerate oxidative and nitrosative stress, suggesting a close interaction between central metabolism, environmental regulation, and genes acquired by horizontal transfer.

Given the ecological and biotechnological relevance of this species, it is imperative to understand the role of ICEs in their genomic evolution. To date, no in-depth study has been conducted on these mobile elements present in this bacterium, analyzing their contribution to it. Nevertheless, the identification and precise classification of ICEs remain challenging due to their modular composition and the prevalence of fragmentation in genomic assemblies (Wang et al., 2024). The present study focused on describing the modular architecture and gene repertoire of these elements, exploring their possible evolutionary origin and phylogenetic relationships with ICEs from other species of the order Sphingomonadales, as well as evaluating their functional potential in the metabolic and ecological adaptation of the *S. granuli* TFA strain, with special emphasis on adaptation modules and genes associated with defense, regulation, and lipid metabolism. When considered as a whole, this analysis provides a more comprehensive perspective on the diversity and evolutionary role of ICEs in catabolically versatile and environmentally relevant bacteria.

## Methods

### Identification of mobile elements in *Sphingopyxis granuli* TFA and conservation analysis

The ICEberg v3.0 tool was utilized to identify integrative and conjugative elements (ICEs). This tool predicts ICEs based on the detection of genes characteristic of integration and conjugative transfer, as well as the presence of regions flanked by specific insertion sequences (Wang et al., 2024). The genome of the *S. granuli* TFA strain (NZ_CP012199.1) was employed as the reference genome. The sequences of the identified ICEs were downloaded and subjected to a homology analysis using BLASTN searches against the NCBI database (Camacho et al., 2009). The objective of this analysis was to identify ICEs present in both genomes of the same species and genomes of related species. A minimum coverage threshold of 60% and an identity of 70% was applied.

### Phylogenetic inference and genomic comparison

The evolutionary relationship between *S. granuli* TFA and the species in which mobile elements were identified was assessed using phylogenetic inference based on the 16S rRNA gene and genomic similarity analysis employing average nucleotide identity (ANI).

The 16S rRNA gene of all analyzed species was initially identified using Barrnap (Seemann, 2013), and its annotation was validated against the SILVA v138.2 database. The sequences were aligned using MAFFT v7.526 with the FFT-NS-2 algorithm, employing the default parameters (Katoh & Standley, 2013). Subsequently, phylogenetic inference was performed using maximum likelihood with the IQ-TREE v3.0 tool and the GTR+I+G substitution model (Minh et al., 2020), which was automatically selected with ModelFinder (Kalyaanamoorthy et al., 2017). The robustness of the nodes was evaluated using 1,000 ultra-fast bootstrap (UFBoot) replicates, and the resulting tree was visualized and annotated in R with ggtree v3.2.1(Yu et al., 2017).

To complement the phylogeny and evaluate the genomic proximity between the analyzed taxa, the Average Nucleotide Identity (ANI) was estimated using skani v0.3.0 (Shaw & Yu, 2023), retaining for each comparison the identity and coverage values calculated by the tool. The generated ANI matrix was graphically represented using a heat map constructed in R with gheatmap v1.4.11.

### Structural and functional characterization of mobile elements

The structural annotation of the identified mobile elements was performed using Bakta v1.11.4, a software that allows for standardized prediction of genes and genomic features through curated models and high-precision ORF detection (Schwengers et al., 2021). The annotations thus generated were then compared with the annotations available in NCBI for the reference genome. This process was undertaken to validate the structural consistency of the predicted genes and to detect possible discrepancies or improvements in the delimitation of reading frames.

To obtain a comprehensive functional characterization, multiple complementary approaches were integrated. First, the encoded proteins were annotated using eggNOG-mapper v5.0.0, assigning COG functional categories and orthologies inferred from phylogenomic databases (Cantalapiedra et al., 2021). Furthermore, annotations were derived from Sma3S v2.0.0, with UniProtKB serving as the reference database to retrieve Gene Ontology (GO) terms associated with biological processes, cellular components, and molecular functions (Bairoch et al., 2005; Casimiro-Soriguer et al., 2017). The identification of protein domains and functional signatures was facilitated by InterProScan v5.76-107.0, a tool that enabled the integration of evidence from multiple databases, including Pfam, SMART, TIGRFAMs, and PROSITE, among others (Jones et al., 2014). The R package gggenes v0.5.1 was employed to visualize the annotated genes.

### Pan-genomic analysis of conserved mobile elements

To characterize the genetic variability of mobile elements conserved in other species, all identified sequences were collected and subjected to uniform structural annotation using Bakta v1.11.4 (Schwengers et al., 2021), ensuring accurate and consistent prediction of open reading frames (ORFs) and coding regions. The pangenome was constructed from the resulting annotation files using the Panaroo tool v1.5.2 (Tonkin-Hill et al., 2020), with the following parameters: -f 0.9 -c 0.9 --len_dif_percent 0.9 --clean-mode sensitive --remove-invalid-genes --merge_paralogs. These settings were selected to impose strict thresholds for similarity, coverage, and length, ensuring the removal of spurious genes and avoiding artificial fragmentation of gene groups. The analysis yielded a robust, refined pangenome, which was subsequently divided into four distinct groups: (1) “core” genes, which are present in all sequences; (2) “soft-core” genes, which are present in the vast majority (95–99%) of sequences; (3) “accessory” genes, which are present only in certain strains; and (4) “unique” genes, which are present in a single sequence.

### Acquisition and Integrated Processing of RNA-seq Datasets

Transcriptomic datasets were retrieved from two previously published studies investigating the physiological responses of *S. granuli* TFA under diverse environmental and stress conditions (de Dios et al., 2020; González-Flores et al., 2019). RNA-seq reads were obtained for the following conditions: (i) exponential growth using tetralin (THN) as the sole carbon source; (ii) exposure to tetralin as an organic solvent for 30 minutes (StressTHN); (iii) induction of nitrosative stress through nitric oxide treatment (StressDETANO); (iv) stationary phase in BHB medium, representing nutrient-limitation stress (stressfest); (v) exponential growth under anaerobic conditions; and (vi) growth on tetralin followed by a 30-minute treatment with BHB. Exponential growth in β-hydroxybutyrate (BHB) served as the basal reference condition.

The raw reads were subsequently subjected to a rigorous quality control process, which entailed the implementation of a series of stringent filters. This process involved the removal of adapter sequences using the BBduk software (BBMap suite v39.52) (Bushnell, 2014). Concurrently, standard quality trimming and the removal of contaminant sequences were performed. Subsequently, the cleaned sequences were aligned against the *S. granuli* TFA reference genome using HISAT2 v2.2.0 (Kim et al., 2019). The generated alignments were subsequently processed to obtain gene-by-gene count matrices using HTSeq-Count from the HTSeq v2.0 tool (Anders et al., 2015).

The differential expression analysis was performed with DESeq2, utilizing its moderate dispersion model and internal normalization based on size factors (Love et al., 2014). This approach is well-suited for addressing variations in sequencing depth between libraries. Given that the data originated from independent studies and potentially exhibited batch effects, an additional homogenization process was implemented prior to DESeq2 to enhance the comparability between samples. The initial step involves the scaling of the data according to the maximum and minimum values within each sample. This is followed by an adjustment of the median of the reads per library. This double procedure smoothed out systematic variations between datasets and reduced the influence of technical differences between studies, facilitating the joint integration of expression profiles prior to statistical modeling and biological interpretation.

## Results

### The genome of *S. granuli* TFA contains ICE elements that are not conserved in other strains of the genus *Sphingopyxis*

Initially, the ICE sequences were sought within the genome of *S. granuli strain TFA* (NZ_CP012199.1) employing the ICEfinder tool. Consequently, eight regions were annotated as putative mobile elements in the genome of strain TFA (Figure 1A), which has a length of 4,679,853 base pairs (bp). Specifically, seven possible ICEs were identified and designated 1 to 7 according to their distribution throughout the genome. Notably, only one Integrative Mobilizable Element (IME) was identified. An IME is a distinct category of mobilizable element that possesses the capacity for excision and integration through its dedicated *oriT* and integrases. However, these elements cannot be transferred autonomously; therefore, they rely on conjugative plasmid conjugation mechanisms or co-resident ICEs for transfer (Guédon et al., 2017).

**Figure 1.**
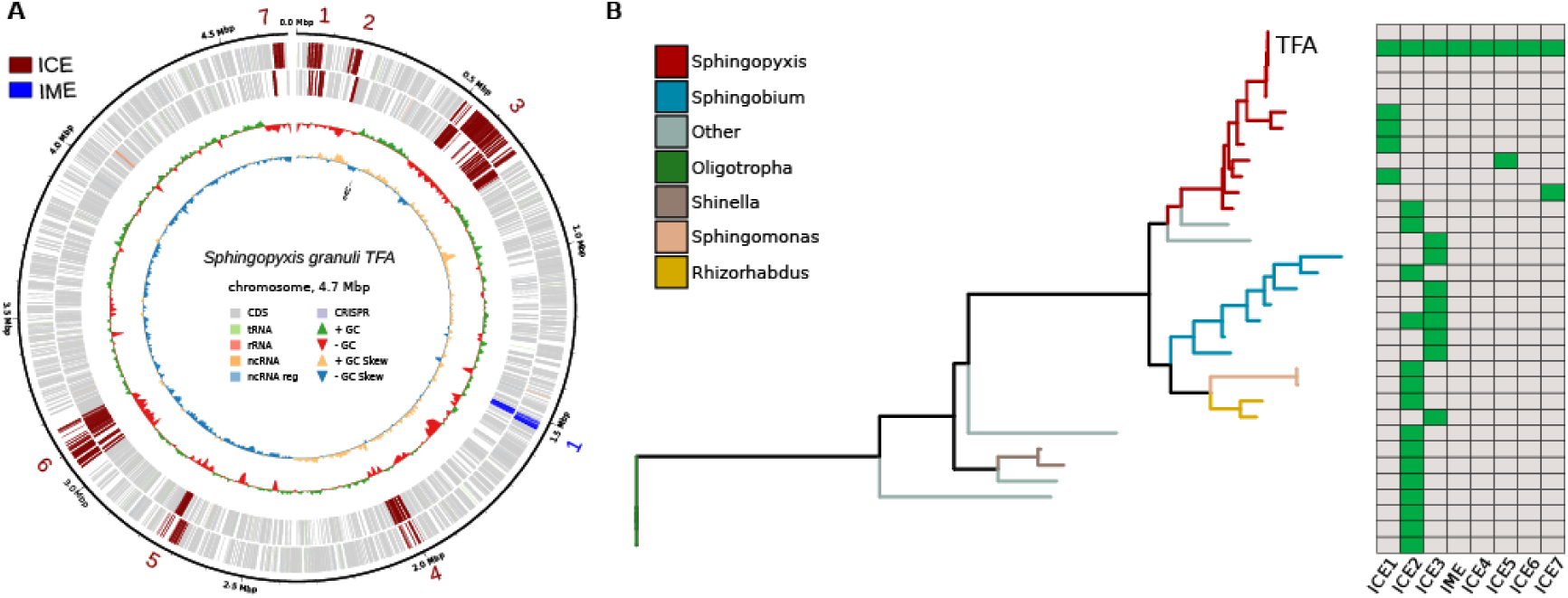
Mobile elements in the genome of *S. granuli TFA* and evolutionary conservation. A) Structural annotation of mobile elements in the genome of *S. granuli TFA*. Mobile elements (red = ICE and blue = IME) and CDS are indicated in different colors. B) Phylogenetic analysis of the 16S sequence of the genomes of *S. granuli* and those with mobile elements shared with the TFA strain. The classification of each clade has been conducted in accordance with the genus of the species. Furthermore, the presence or absence of the various ICEs is indicated by green, which signifies their presence, and gray, which signifies their absence.

To determine the prevalence of ICE sequences in other genomes of the species, a comprehensive analysis of available strains of *S. granuli* was conducted, enabling the calculation of the prevalence of mobile elements in the species. Consequently, it was observed that a limited number of strains possessed mobile elements, attributable to the substantial fragmentation of their genomes (**see Table X**). Furthermore, it was ascertained that only a subset of ICEs from the TFA strain were retained in the remaining genomes (namely, ICE1, ICE5, and ICE7).

Given the propensity of mobile elements, such as ICEs and IMEs, to integrate into other genomes with relative ease, a homology search was conducted on the regions detected in the TFA strain genome across diverse genomes from different evolutionary taxa. Consequently, two ICEs, ICE2 and ICE3, were found to be distributed in other taxonomic groups. However, as previously mentioned, these mobile elements did not appear to be integrated into the other strains of the species *S. granuli*.

The genomes exhibiting these ICEs belonged phylogenetically to α-proteobacteria (NCBI:txid28211), specifically to the order Sphingomonadales (NCBI:txid204457), the same as *S. granuli*, and to the order Rhizobiales (NCBI:txid356), also known as Hyphomicrobiales (Figure 1B). In these genomes, the ICE sequences invariably exhibited exceptionally high sequence identity, sometimes exceeding 90% identity, and optimal average coverage.

In order to verify that only ICE2 and ICE3 were preserved and that this was not an annotation or labelling error in the sequences, the average nucleotide identity (ANI) was calculated between the genomes that had these mobile elements integrated and the strains of the species S. granuli. Initially, it was demonstrated that the *S. granuli* genomes exhibited a high degree of similarity, with more than 97% identity and an average coverage of 75% (Figure 2). In contrast, the remaining additional genomes, which did not belong to the species, had coverages of less than 30% and identities of no more than 90%. Consequently, it was ascertained that solely mobile elements were conserved within these genomes and TFA strain, a result of the low shared coverage obtained. It is important to acknowledge that certain genomes exhibited significant fragmentation, a consequence of metagenomic assembly techniques, which have been shown to specifically modify sequence coverage.

**Figure 2.**
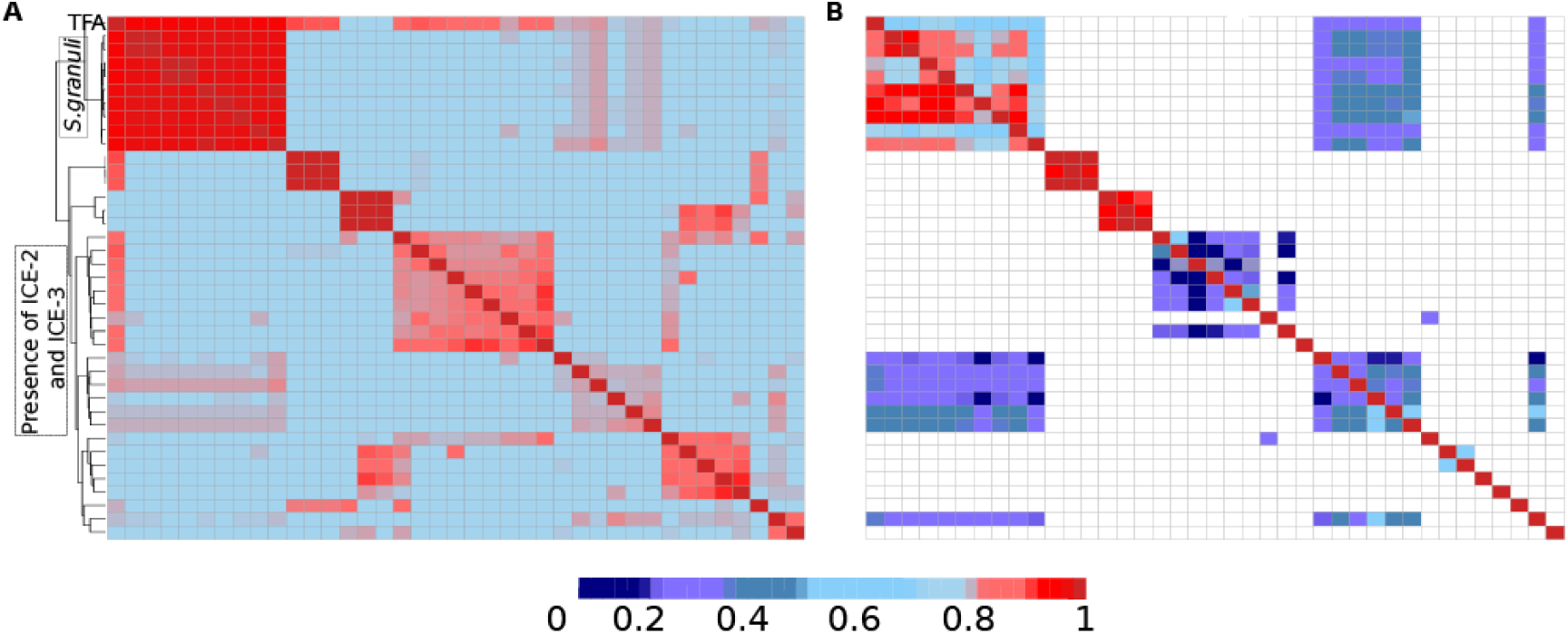
Heatmap representing the results obtained from the ANI calculation. A) The percentage of shared identity. B) The percentage of shared coverage. The strains corresponding to *S. granuli* and those that share at least ICE2 or ICE3 are indicated.

### Most of the identified ICE did not present all of their functional modules

In synthesis, an ICE necessitates the integration/excision module, the conjugation module, and the regulation module. Additionally, the presence of an adaptation module might facilitate its persistence within bacterial populations. Consequently, to assess the functionality of the aforementioned ICEs, the presence or absence of these genomic modules was analyzed.

As a result, it was observed that all these regions had the necessary elements to carry out a conjugation process, since genes of the type IV secretion system (T4SS), coupling genes (T4CP), and a functional relaxase were found in all of them. The majority of these relaxases were of the P type, which is the most prevalent family in plasmids and integrons (Garcillán-Barcia et al., 2009). However, with the exception of the ICE3 and ICE6 regions, these genetic elements did not exhibit the presence of integration or regulation modules, as they lacked their own integrases and regulatory genes. This was primarily due to the fact that these ICEs (ICE1, ICE2, ICE4, ICE5, and ICE7) had an average size of less than 20 kb, which differed from the size of the other two ICEs, 218 kb and 81 kb, respectively. It is noteworthy that a significant number of functional integrase genes were identified in these larger ICEs (Figure 3).

**Figure 3.**
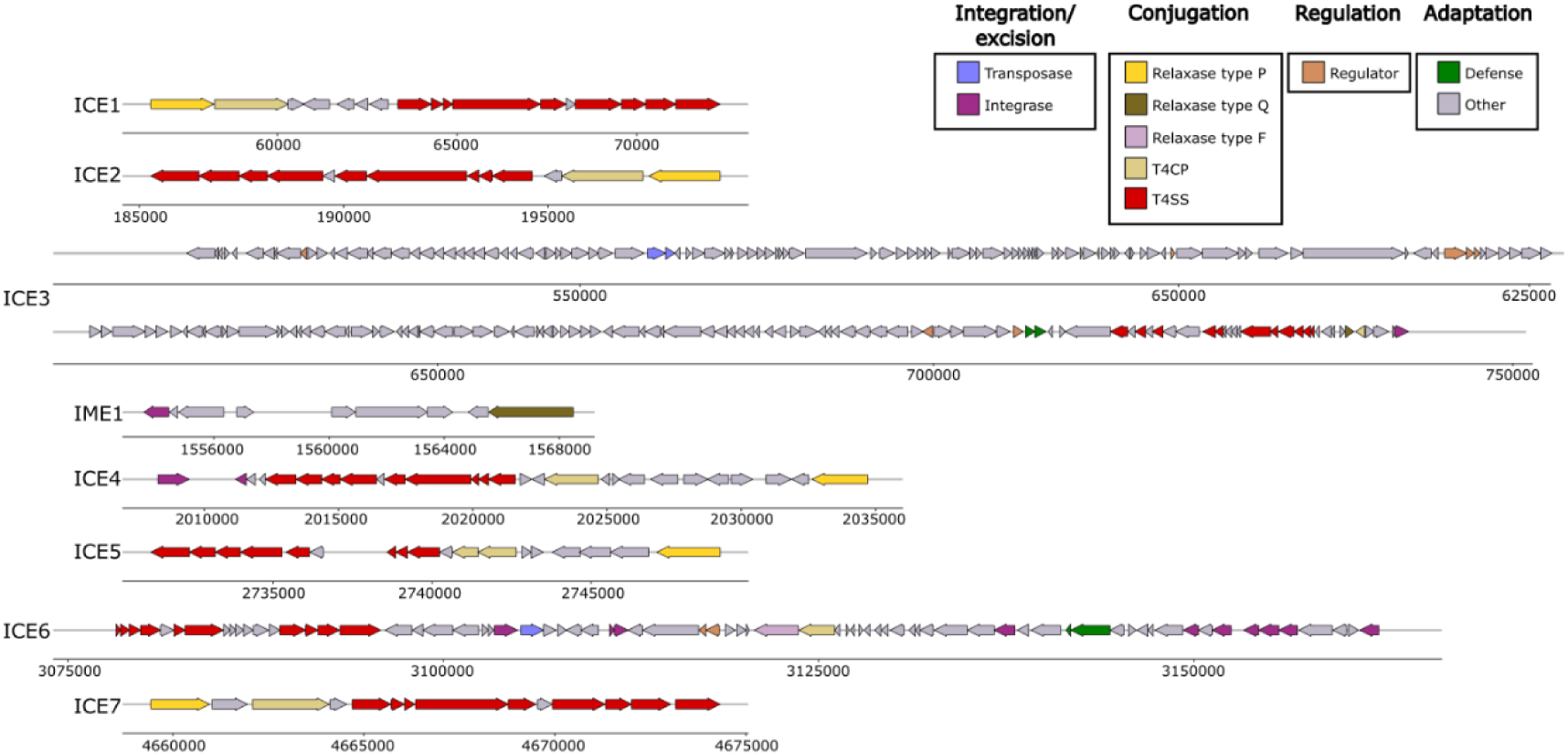
Modular organization of the ICEs analyzed. The figure illustrates the modular organization of the ICEs under analysis, emphasizing the primary functional components: integration/excision (integrase and transposase), conjugation (relaxase, type IV secretion system (T4SS) proteins, and T4CP coupling protein), adaptation modules (defense systems and other genes), and potential regulatory modules (MarR families, response regulator, among others). Each module is represented by the characteristic genes identified in the analysis.

The ICE3 analysis identified several genes with domains associated with transcriptional regulation and adaptive control (Figure 3). These include a member of the MarR family (*MarR_2*), several regulators with helix–turn–helix motifs (HTH proteins, RHH proteins, *TetR_N*, and *GerE*), and components characteristic of two-component systems (*parA* and *parB*). In contrasts, the regulatory module in ICE6 was predominantly comprised of genes encoding WYL (Trp–Tyr–Leu) domain-containing transcriptional regulators and an S24 protease. These regulators modulate the expression of genes related to conjugation and transfer depending on external or physiological signals (Makarova et al., 2014).

To conclude the analysis of the ICE modules themselves, several genes encoding proteins were identified which, based on their known functions, could form a possible adaptation module. In accordance with the greater total length of ICE3, its adaptation module was found to be more extensive than that of ICE6. Despite this variation in size, genes associated with bacterial defense mechanisms were identified in both ICEs, although with differences in the systems present: ICE3 contains genes corresponding to the AbiE system, which is related to defense against phages through infection abortion mechanisms (Dy et al., 2014). ICE6, meanwhile, carries genes from the Hachiman system, which was recently described as a broad-spectrum bacterial immunity system (Millman et al., 2022). Furthermore, restriction-methylation genes were identified in both ICEs, indicating a potential complementary strategy to evade host defense systems and facilitate the establishment and transmission of the element. It is noteworthy that a significant proportion of the genes identified in these modules are hypothetical proteins, accounting for nearly 70% of the total. This observation poses a limitation on current functional inferences and underscores the necessity for more comprehensive functional annotation.

### Functional annotation reveals a large number of genes associated with accessory metabolism and resistance in the ICE3 region

To provide a thorough characterization of the adaptive module of the ICEs under analysis, a systematic annotation of all their coding genes was conducted. To this end, prediction and functional assignment tools widely used in comparative genomics, such as InterProScan, Sma3s and eggNOG-mapper, were employed to identify conserved domains, protein families, and possible biological functions from 213 and 73 amino acid sequences of the ICE3 and ICE6 regions, respectively. Consequently, there was a substantial decrease in the proportion of hypothetical genes, that is, genes for which no known function has been determined. In ICE3, this percentage underwent a significant shift, increasing from an estimated 67% in previous annotations to 20% in this study. Similarly, in ICE6, the percentage decreased to approximately 16%, reflecting a high degree of functional resolution. Additionally, tRNA was predicted, as these are sites of ICE insertion. It was observed that only ICE3 had direct repeats flanking it. In contrast, ICE6 only showed a tRNA-lys (Figure 4).

**Figure 4.**
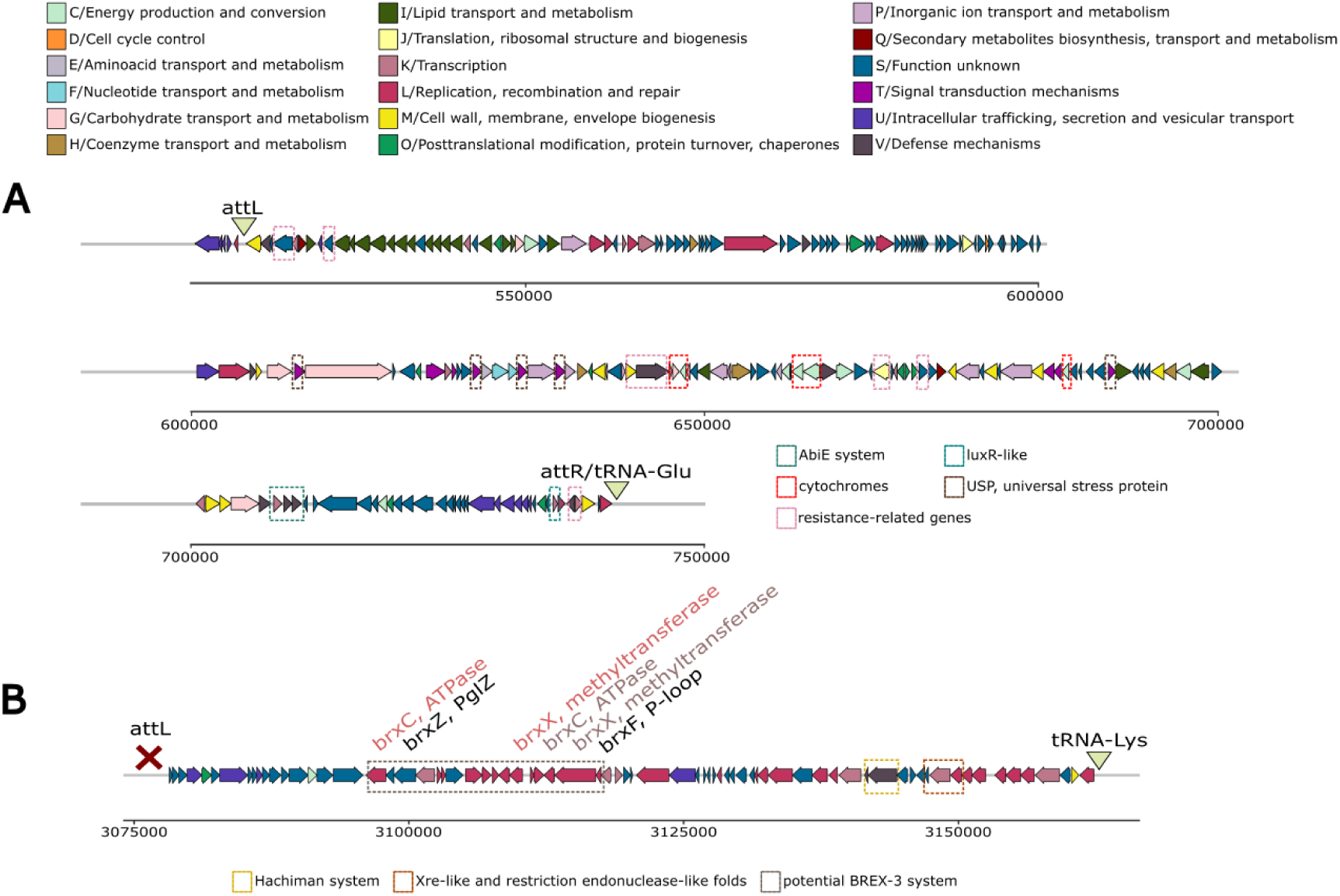
Functional annotation of the ICE3 and ICE6 mobile elements. A) Functional annotation of ICE3. B) Functional annotation of ICE6. Genomic elements characteristic of each ICE has been designated with different colors (dashed squares). Each gene is represented by a block colored according to its functional category, as assigned by the COG (Clusters of Orthologous Groups) system. This category was obtained using eggNOG-mapper. The categories encompass functions such as metabolism, transcription, replication, cell cycle control, and unknown functions, among others. In the BREX-3 system, components have been identified with different colors. The functional distribution enables the visualization of the adaptive and regulatory potential of each ICE, as well as the identification of regions enriched in specific functions. Note that ICE6 did not present the direct insertions characteristic of an ICE.

As previously mentioned, the mobile elements in question exhibited a typical modular architecture, with genes associated with integration and excision, transfer via type IV secretion systems (T4SS), regulation, maintenance, resistance, and a wide range of accessory functions. It is noteworthy that in ICE3, a gene was identified that encoded an accessory protein, MobA, which was necessary for processing *oriT* by Q-type relaxase, as present in this ICE (Garcillán-Barcia et al., 2019).

With regard to the regulatory module, novel potential components were identified in both ICEs. In the case of ICE3, a gene encoding a LuxR-like protein was identified. This family of proteins has been described as sensor-regulators in bacteria (Subramoni & Venturi, 2009). Additionally, it emphasized the presence of a VapB/VapC system, which is known to stabilize mobile elements, thereby promoting their retention in the bacterial population through the negative selection of cells that lose these elements (Hollingshead et al., 2024). In contrast to previous findings, the analysis performed in ICE6 revealed the presence of two *copG* and *Xre-like* genes. These genes, classified as transcriptional repressors, have been characterized in mobile elements such as plasmids and phages (Hernández-Arriaga et al., 2009; Long et al., 2022).

It is noteworthy that the adaptive module played a particularly salient role in ICE3 (Figure 4A). A substantial number of genes implicated in the degradation of lipids (e.g., acyl-CoA dehydrogenase or lipase/esterase), carbohydrates (e.g., β-glycosidase), and aromatic compounds (e.g., mono- and dioxygenases) were identified in this mobile element. ICE3 carried genes encoding cytochromes, β-lactamases, and multiple USP (universal stress protein) genes in its region. In addition to these genes, other elements encoding key resistance determinants were annotated, such as RND-type efflux pumps, MFS transporters, chromium detoxification genes or AbiE system (Figure 4A).

In ICE6, while the accessory metabolic repertoire was more limited, genes with restriction endonuclease-like folds associated with stress response (RES) domains and strawberry notch-type helicases associated with regulatory functions and chromatin remodeling were identified. Consequently, this mobile element presented metalloproteases, additional anti-restriction factors, and components of the BREX-3-type bacterial defense system. Initially, the system appeared to be incomplete, containing only the *brxZ* (protein encoded with PglZ domain) and *brxF* (P-loop NTPase) genes. However, subsequent to the association of classic components with the annotation of several nearby genes in the same orientation, the defense system was completed with the *brxC* (ATPase) and *brxX* (methyltransferase) genes. It is important to note that this defense system could be completed even though it is fragmented by genes related to transposases or integrases (Figure 4B).

### The ICE3 pangenome reveals a dual architecture shaped by integration-proximal cores and distal variable hotspots

Given the conservation of the ICE3 element in other species that do not belong to the *Sphingopyxis* genus, a conservation analysis of this element was conducted. To this end, a “pangenome” of the region comprising this ICE was constructed, with the aim of identifying the genetic variability that this sequence could have. The term “pangenome” is employed to denote the complete set of genes present within a given species (Tettelin et al., 2005). In this context, the species of interest is the mobile sequence ICE3. In this pangenome, the core genes, which are essential genes shared by all ICEs, and the accessory genes, which are present in only some ICEs, were analyzed. The construction of this pangenome was achieved through the selection of all ICE3 sequences located in the initial section of results. These sequences were subsequently annotated, and the resulting genes were grouped according to identity and coverage thresholds, thereby creating robust gene clusters.

The resultant initial pangenome comprised over 491 gene clusters, of which only 16 genes were present in all ICE sequences across the various species. These genes were associated with universal stress proteins and transferases involved in the metabolism of diverse compounds. Despite the high average identity detected, some sequences exhibited low coverage with ICE3. To verify this, we assessed the presence or absence of each cluster in the different strains. The analysis revealed that two sequences, from *S. phenoxybenzoativorans* and *R. wittichii*, shared very few pangenome clusters, as they formed a large number of independent clusters containing their own genes. Furthermore, the ICE from *Sphingobium sp. MI1205* showed minimal sequence coverage compared to the other sequences, despite lacking exclusive clusters (Figure 5A). These three ICE sequences were excluded from subsequent pangenomic analyses to obtain the most complete pangenome possible.

**Figure 5.**
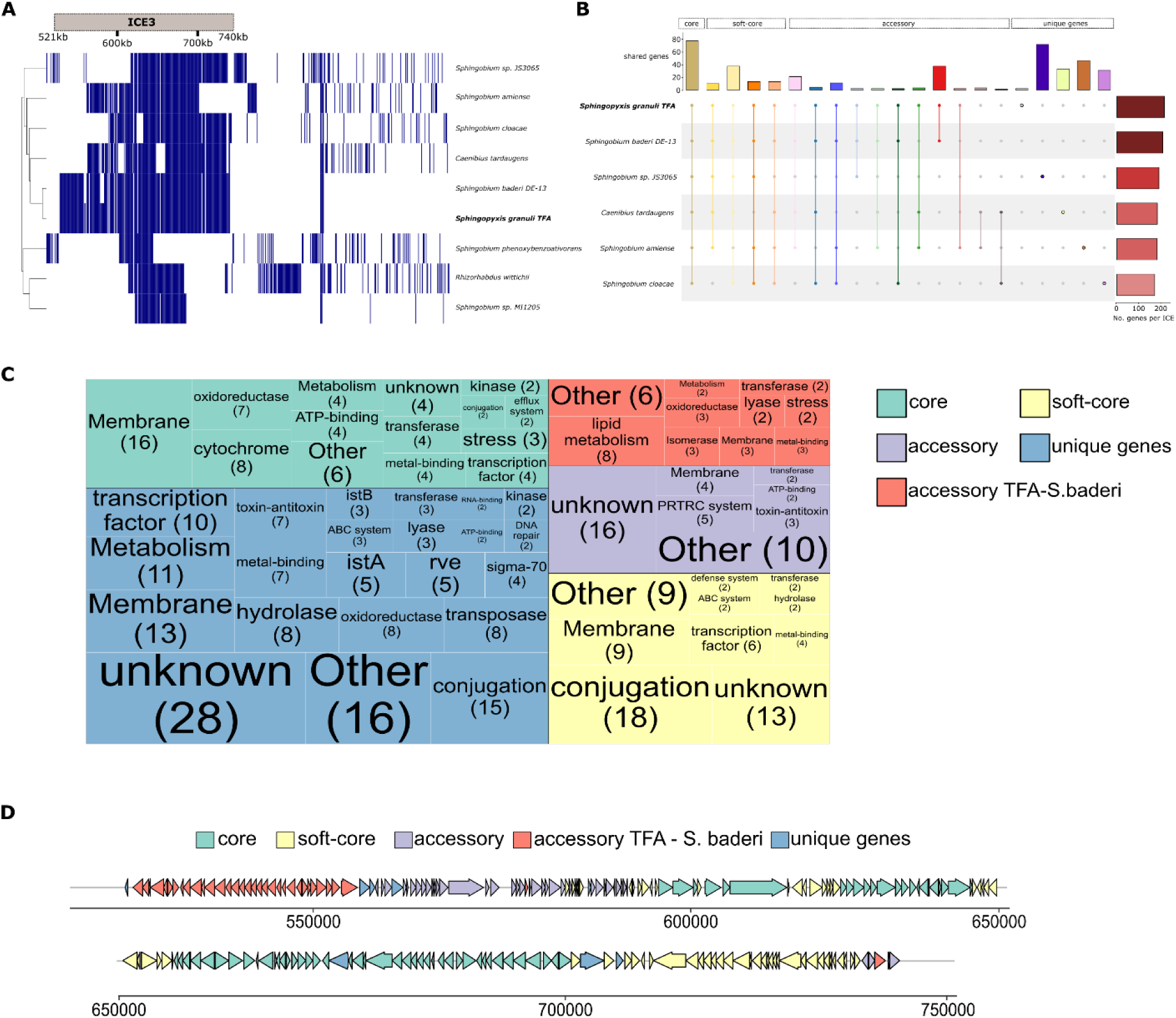
Analysis of the ICE3 pangenome. A) Presence/absence of the genes that constitute the pangenome. The genes were ordered with respect to the species *S. granuli* TFA, indicating the presence of the gene in blue. B) Upset plot of the pangenome. Count of the number of genes according to pangenome group and distribution according to ICE3 variant in each species. C) Treemap of functional terms of the pangenome. Each rectangle represents a functional category, where the size reflects the number of genes with that functional annotation. They have been marked with different colors according to pangenome group. D) Distribution of pangenome groups in the ICE3 structure.

Subsequent to the calculation of the new pangenome, a total of 386 genes were identified, of which 70 were present in all sequences, representing a considerable increase compared to the initial pangenome. In contrast, soft-core and accessory genes were distributed in smaller proportions. A significant finding was the presence of a large number of unique or exclusive genes within the various ICEs (with an average of 30 to 40 genes), with the ICE found in the species *Sphingobium* sp. JS3065 containing the highest number of exclusive genes (65). Regarding the species *S. granuli* TFA and *Sphingobium baderi* DE-13, numerous clusters exclusive to both species were identified, demonstrating their high sequence similarity and coverage (34 shared genes) (Figure 5B). These different categories of the pangenome were then subjected to functional analysis in order to assess the essential and accessory functions acquired by the distinct ICE contexts across different species.

Comparative analysis of pangenome categories revealed pronounced differences in the functional composition of ICE core and soft-core genes (Figure 5C). The core repertoire was dominated by functions essential for element stability and basal transfer, including relaxase, the TraD coupling protein, and channel components (*tolC*, *oprM*), together with resistance determinants (*arsR*, *czcA/B*) and universal stress response factors (usp family, *ahpC, trxC, dps*). By contrast, the soft-core set was enriched in *tra/trb* genes encoding the complete type IV transfer machinery. It also incorporated diverse transcriptional regulators and two-component systems (*fixL/fixJ, CopG, LuxR/XRE*), as well as antibiotic resistance (*aph*), stress response (*hspC2*), and metabolic functions (*sulP, phnA, bktB, pckA*). Collectively, these findings indicate that the core constitutes the minimal backbone required for ICE maintenance and transfer, whereas the soft core confers modularity and plasticity, facilitating adaptation across bacterial hosts and environmental niches.

Accessory genes included multiple functions related to bacterial resistance and defense, such as components of MFP-type efflux pumps and VapB/VapC toxin–antitoxin systems, as well as elements involved in genetic regulation and stability (*parB*). Furthermore, it is essential to emphasize the presence of a complete Pilus Retraction/Contraction (PRTRC) System related to the formation and dynamics of type IV pili in bacteria (Figure 6C). Accessory genes exclusive to the mobile elements of *S. granuli* TFA and *S. baderi* were analysed separately, revealing numerous transport- and secretion-related functions (*nodT*, *ydhJ*, among others). In addition, several MarR family transcriptional regulators were detected, together with genes implicated in the degradation and biosynthesis of lipids, fatty acids, and aromatic compounds.

**Figure 6.**
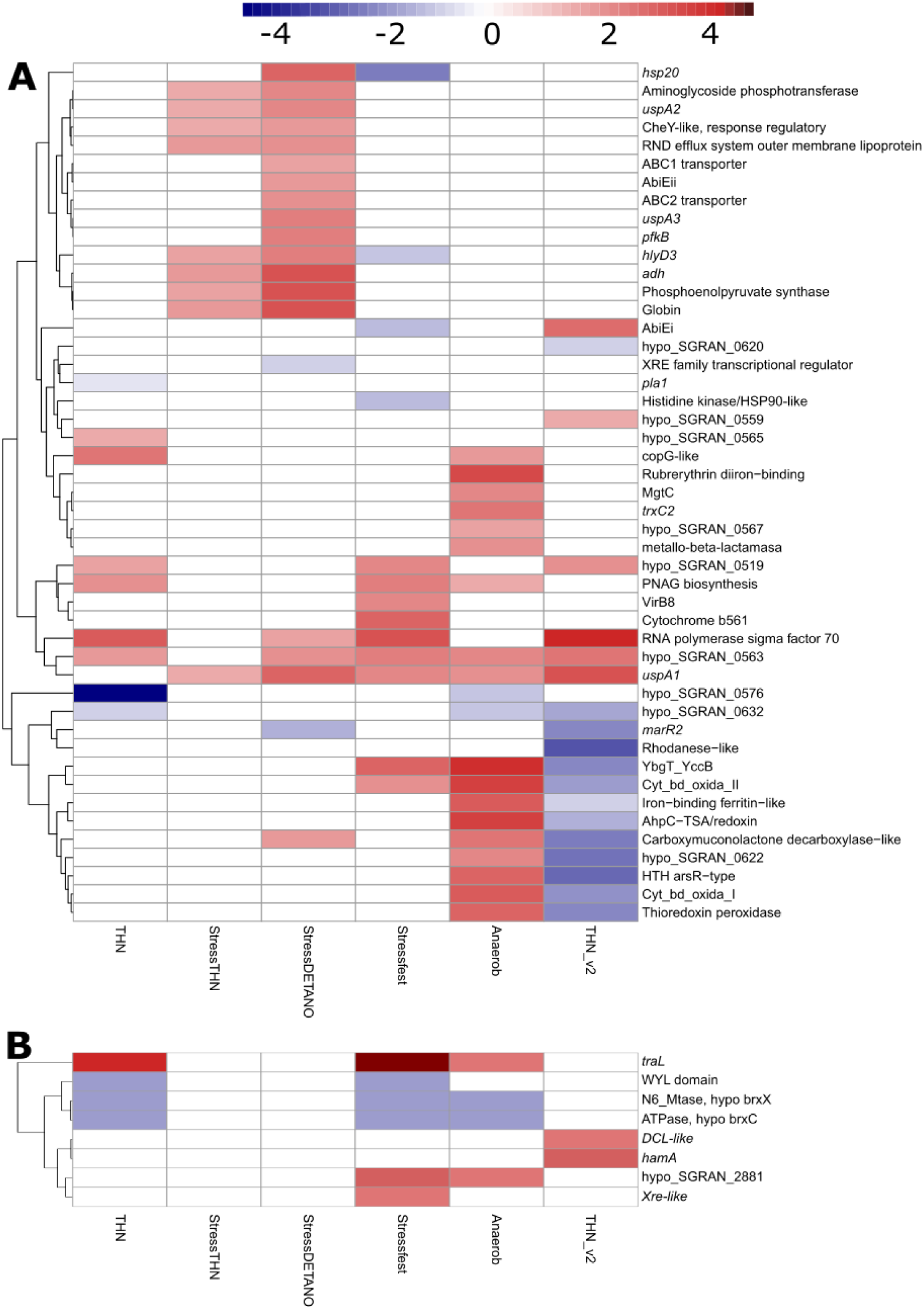
Heatmap of differential gene expressions under six experimental conditions. A) A heatmap illustrating the differential regulation of genes from ICE3. B) A heatmap depicting the differential regulation of genes from ICE6 under six experimental conditions. Genes that are overexpressed (upregulated) or repressed (downregulated) compared to the baseline condition are shown. The colors of the markers indicate the type and magnitude of regulation (log fold change values), allowing specific patterns of activation or silencing to be visualized (red = overexpression, blue = repression). The conditions evaluated include: (1) exponential growth in tetralin as the sole carbon source (THN), (2) acute exposure to tetralin for 30 minutes in culture with BHB (StressTHN), (3) nitrosative stress induced by nitric oxide (StressDETANO), (4) stationary phase in BHB medium (Stressfest), (5) anaerobic growth in exponential phase, and (6) culture in tetralin with acute exposure to BHB for 30 minutes (THN_v2). The ICE3 region exhibits a higher proportion of regulated genes, while ICE6 demonstrates limited differential expression, suggesting more restricted activation dependent on specific signals.

To conclude the functional analysis, exclusive genes were examined, revealing a large repertoire of transposases from diverse families. Additional resistance and tolerance determinants were also detected, including the *cusC* cation efflux system, multifunctional ABC transporters, and sigma/antisigma factors. These results suggest that the acquisition of specific gene variants may occur with relative ease. Notably, the ICE in *Sphingobium* sp. JS3065 carried a distinct conjugation system together with genes involved in morphogenesis and cell envelope remodeling, including *mltC, mrdA, mrdB, mreB, mreC,* and *mreD*, which are central to peptidoglycan biosynthesis and cell shape control.

The distribution of pangenome groups within the ICEs revealed a clear non-random organization (Figure 5D). Utilizing the ICE of *S. granuli* TFA as a reference point, a comparative analysis was conducted of the positioning of conserved genes across the various elements. Core and soft-core genes were predominantly clustered at the initial region of the ICE, in close proximity to the tRNA integration site, underlining the strong conservation of this portion of the element. In contrast, as the distance from the integration point increased, the proportion of accessory and species-specific genes became markedly higher, particularly in the ICEs of *S. granuli* TFA and *S. baderi*.

Overall, the observed distribution supports a modular architecture in which the conserved regions located near the tRNA integration site safeguard the essential functions of the element, while the peripheral, more variable regions provide evolutionary plasticity. This spatial arrangement may facilitate the balance between conservation of the ICE backbone and diversification through the acquisition of new genes, thereby enhancing the adaptability of these mobile elements in different bacterial hosts and environmental contexts.

### ICE3 and ICE6 have been identified as pivotal architects of bacterial resilience to stress conditions

In order to analyze the importance of the gene content of ICE, ICE3, and ICE6, a comparative transcriptomic study was performed using RNA-seq. To this end, the exponential growth of *S. granuli* TFA with β-hydroxybutyrate (BHB), a simple and directly assimilable carbon source, was established as the baseline condition. According to this reference, an evaluation of various physiological and environmental stress scenarios was conducted, with the consideration of a normalization process that mitigated a hypothetical RNA-seq batch effect.

First, exponential growth was analyzed using tetralin (THN) as the exclusive carbon source, thereby representing a more advanced metabolic scenario. Complementarily, an acute exposure condition was included in which exponentially growing cells with BHB were treated for 30 minutes with tetralin (StressTHN). To further investigate an alternative form of cellular stress, the effects of nitric oxide, acting as a nitrosative stress agent (StressDETANO), were examined. Similarly, the stationary phase in BHB (Stressfest) was selected as a model of stress associated with nutrient limitation. Conversely, the transcriptomic response under anaerobic conditions in the exponential phase was analyzed, with the aim of evaluating the impact of oxygen deprivation on central metabolism (Anaerob). Finally, the condition inverse to the StressTHN treatment was analyzed, in which cells growing in tetralin were exposed for 30 minutes to BHB (THN_v2).

In the case of ICE3, of the 213 genes constituting the region, 47 genes demonstrated at least one positive/negative deregulation in some experimental condition studied (Figure 6). In the THN condition, robust induction of the sigma-70 factor and a *copG-like* gene was observed, along with hypothetical genes such as SGRAN_0563 and SGRAN_0519, suggesting partial reprogramming associated with hydrocarbon metabolism. In the StressTHN condition (acute exposure to tetralin), genes associated with transport and resistance, such as *uspA2*, aminoglycoside phosphotransferase, and *hlyD3*, were activated. However, no significant negative alterations were observed among the top regulated genes, suggesting a swift response to the xenobiotic.

A considerable increase in the number of activated genes was observed under nitrosative stress (NO). Consequently, antioxidant genes such as globin, *AhpC-TSA/redoxin*, and thioredoxin peroxidase were induced, suggesting an adaptation to reactive nitrogen species. In contrast, only two genes associated with genomic regulation, *marR2* and *XRE-like*, underwent gene repression. During the stationary phase in BHB, a marked induction of the sigma factor and several genes related to cytochromes was observed again, reflecting an adaptation to reactive nitrogen species and metabolic reorganization. Under this condition, a clear deregulation of *hsp* genes was obtained.

The anaerobic condition exhibited a marked induction of genes associated with oxidative stress, including rubrerythrin, ferritin, *trxC2*, and various cytochromes. This induction suggests a reconfiguration of redox metabolism. Finally, in the THN_v2 condition (BHB shoot in tetralin), a clear deregulation of cytochromes and genes associated with iron regulation was observed, indicating a metabolic transition towards the use of BHB.

It is noteworthy that several genes exhibited differential regulation under multiple conditions, including abiE system, *uspA1*, hypo_SGRAN_0563, and sigma-70 factor. This finding suggests consistent and recurrent responses of ICE3 to diverse stresses.

Transcriptomic analysis of the mobile element ICE6 revealed limited differential regulation, with only a small subset of genes showing significant changes in expression, even though ICE contains a total of 73 genes. This low proportion of regulated genes suggests highly controlled expression, possibly conditioned by specific stimuli. It is noteworthy that three genes related to a possible BREX-3 system exhibited a highly analogous expression pattern, being repressed under THN, stationary phase, and anaerobic conditions.

In the stationary phase, a notable induction of *traL* was observed, accompanied by the expression of hypothetical genes such as hypo_SGRAN_2881 and the *Xre-like* regulator. This suggests the potential activation of functions related to genetic transfer and metabolic adaptation. Except for *Xre-like* genes, the remaining genes demonstrated expression under anaerobic conditions. Finally, under the THN_v2 condition, genes such as *hamA* and *DCL-like* were activated, possibly involved in signaling or structural remodeling.

This experimental design enabled a comparative analysis of baseline conditions with various types of metabolic, oxidative, nitrosative, and physiological stress. It thereby established a comprehensive framework for the analysis of transcriptomic plasticity provided by both ICEs.

## Discussion

This study has thoroughly characterized the structure, evolutionary conservation, functional organization, and gene expression of integrative and conjugative elements (ICEs) identified in the genome of *Sphingopyxis granuli* TFA. To this end, evolutionary approaches, comparative annotation, and transcriptomic analysis under multiple physiological and stress conditions have been combined. The genus *Sphingopyxis* comprises gram-negative bacteria that are distributed widely in terrestrial and aquatic environments. These bacteria are distinguished by their remarkable metabolic capacity to degrade aromatic compounds and complex organic pollutants (Sharma et al., 2021). In particular, the *S. granuli TFA* strain has been shown to possess the capacity for the degradation of complex aromatic compounds, with a particular emphasis on polycyclic hydrocarbons and petroleum derivatives. This process utilizes tetralin, also known as 1,2,3,4-tetrahydronaphthalene, as the exclusive source of carbon and energy for its metabolic activities (García-Romero et al., 2020b). In this context, mobile genetic elements such as ICEs play a fundamental role in the ecological adaptability and expansion of the catabolic repertoire of these bacteria, facilitating the acquisition of genes involved in the degradation of xenobiotics, resistance to environmental stresses, or defense against antimicrobial agents (De Maayer et al., 2015; Guédon et al., 2025; van Vliet et al., 2021).

The genomic analysis of S. granuli TFA facilitated the identification of six integrative and conjugative elements (ICEs) through the utilization of specialized bioinformatics tools. These tools employ predictive models based on genetic signatures associated with integration, conjugation, and maintenance processes (Wang et al., 2024). Of these six elements, only ICE2 and ICE3 exhibited conserved regions in other species of the order Sphingomonadales, suggesting that both constitute functionally relevant and evolutionary stable genetic modules within this bacterial group. The preservation of ICE2 and ICE3 in species belonging to the same taxonomic order can be attributed to their direct involvement in metabolic and ecological adaptation to their respective environments. This type of symbiotic association between ICEs and specific catabolic pathways has been previously been documented in environmental bacteria, where mobile elements act as evolutionary vehicles for the transfer of degradation or resistance genes (de Assis et al., 2022), and even for the active defense of the bacterium against phages (Johnson et al., 2022). The preservation of these sequences across disparate taxa can result in annotation inaccuracies, such as contaminated regions that are, in fact, derived from horizontal transfer. This is due to both processes introducing foreign DNA into genomes (Capra et al., 2013). Therefore, novel algorithms were developed to enable the differentiation of these regions (Bálint et al., 2024).

These types of mobile elements are characterized by their high degree of modularity and mosaic nature, arising from recurrent recombination processes and gene acquisition through horizontal transfer. This phenomenon can result in the loss or replacement of essential modules, which, in turn, can lead to partial or ambiguous detection using bioinformatics tools (Wozniak & Waldor, 2010b). A notable caveat is the frequent ambiguity in the delineation of integration boundaries. This is primarily attributable to two factors: first, the non-conservation of all sites; and second, the inherent variability of repetitive or intergenic regions, both of which pose challenges to the assembly process. For these reasons, this phenomenon is exacerbated in genomes that are not completely closed, such as those used in most of this study, where fragmented contigs prevent the complete architecture of the element from being reconstructed, underrepresenting the total length identified in these mobile elements (Cury et al., 2017).

The majority of identified ICEs, with the exception of ICE3 and ICE6, exhibited characteristics indicative of residual presence, exhibiting a lack of functional modules essential for the regulation and integration of these genomic elements. The absence of these modules gives rise to two possible hypotheses. First, these regions are non-functional and have become fixed in the bacterial chromosome as markers of different horizontal transfer events. This has been described in various genera of Alphaproteobacteria and Sphingomonadales. Or, conversely, they are parasitic ICEs that depend on the machinery of other systems, as they are capable of using homologous recombination systems or integrase encoded in other mobile elements or even prophages (Bellanger et al., 2014). This mechanism has been previously described in the Actinobacteria class, where it involves the use of DDE recombinases or specialized relaxases capable of mediating integration/splicing (Ghinet et al., 2011). Consequently, these small ICEs might be non-autonomous and contingent on external elements yet remain functional. These regions can be regarded as hotspots for the integration of genes that confer an adaptive advantage to the bacterium. However, it is noteworthy that these regions did not exhibit a high number of genes in the adaptive module.

The prevalence of genes with unknown functions within adaptation modules is a recurring characteristic of ICEs from environmental bacteria, suggesting a potential for functional diversification that remains to be thoroughly investigated (Che et al., 2024; Meygret et al., 2019). In the mobile elements analyzed in *S. granuli* TFA, the percentage reached 80%. Consequently, a substantial enhancement was achieved in the functional annotation of ICE3 and ICE6 identified in *S. granuli* TFA, enabling a comprehensive analysis of the adaptation module.

The identification of genes associated with bacterial defense systems, including Hachiman, BREX-3 (Bacteriophage Exclusion), and XRE-like regulators, in ICE6 suggests that this mobile element plays a role in both gene dissemination and host protection and stability. The Hachiman system, recently described as a component of bacterial defense mechanisms, has been associated with resistance to phage infections and the restriction of exogenous DNA. However, the precise molecular mechanism underlying these processes remains to be fully elucidated (Doron et al., 2018). Complementarily, BREX systems, and particularly the BREX-3 subtype, have been shown to mediate an epigenetic form of defense that prevents phage replication without degrading their DNA, thus contributing to a non-destructive barrier against viral invasion (Drobiazko et al., 2025). In contrast, XRE-like regulators are often linked to transcriptional control networks that regulate the expression of genes associated with defense, stress, and genetic mobility, as toxin-antitoxin systems and conjugative operons (Long et al., 2022). The coexistence of these three systems in ICE6 suggests a coordinated defense and self-regulation strategy, through which ICE could increase the host’s tolerance to external agents and preserve its own genetic integrity.

In contrast, ICE3 is distinguished by its gene content, which is oriented towards stress response and metabolic adaptation. The presence of multiple USP (Universal Stress Protein) proteins, which are involved in resistance to oxidative conditions and energy limitations, is particularly notable (Luo et al., 2023). Additionally, the existence of genes for cytochromes and lipid metabolism enzymes suggests a functional integration with the host’s respiratory network. Furthermore, ICE3 contains an Abi (Abortive Infection) system, which is often linked to antiviral defense mechanisms that involve the interruption of the phage infection cycle (Dy et al., 2014). This system could confer a selective advantage in environments with high phage pressure, thereby reinforcing the role of ICE3 as a cellular resilience module. This phenomenon has been previously documented in other ICEs (Johnson et al., 2022). The coexistence of genes associated with defense, metabolism, and stress indicates that ICE3 may serve as a multifunctional adaptation element, thereby enhancing the survival of *S. granuli* TFA in dynamic and contaminated environments.

Overall, the findings imply that, while ICE6 primarily functions in the context of genetic stability and defense against exogenous elements, ICE3 may be more directly implicated in physiological plasticity and adaptive responses to the environment. This suggests the potential for ICE3 to serve as a multifunctional module of cellular resilience. The observed disparities in the adaptation modules of ICE3 and ICE6 imply that each element has developed distinct strategies to optimize its persistence and transmission in varied bacterial contexts. The observation that ICE3 possesses a broader adaptation module is consistent with its larger overall size, which could afford it a greater diversity of accessory functions. However, the presence of restriction-modification systems in both ICEs lends further support to the hypothesis that these elements not only depend on the conjugation machinery for their propagation, but also integrate active defense modules that ensure their maintenance in new hosts.

The conservation of ICE3 in other species has enabled the creation of a pangenome of this mobile element. In this pangenome, it was found that the genes in proximity to the integration site, which are typically associated with tRNA genes, exhibited conservation across all sequences. In contrast, the more distal regions demonstrated significant genetic variability. This configuration is indicative of the characteristics of this particular type of mobile element, as it reflects their modular organization and their specialization for the acquisition of new accessory components (Wozniak & Waldor, 2010b). In this distribution, core genes are typically implicated in processes such as integration, excision, and transfer, thereby ensuring the stability and functionality of the element. Concurrently, accessory regions, situated towards the distal ends, concentrate adaptive genes associated with resistance, metabolism, or stress response. This spatial configuration promotes the preservation of functions critical for horizontal mobility while concurrently enabling evolutionary diversification through the acquisition or loss of adaptive modules (Touchon et al., 2014).

In the case of ICE3, the core module also included transposase genes and stress response proteins, such as universal stress proteins (USPs), suggesting that the basic machinery of these elements not only ensures their mobility but also their ability to respond to adverse environmental conditions (Luo et al., 2023). These “basic functional” genes have been previously described in other ICE systems, where it has been proposed that they reinforce the stability of the element and the survival of the host during processes of horizontal transfer or metabolic stress (Johnson & Grossman, 2015). Pangenome analysis revealed a limited number of shared accessory genes, accompanied by a high proportion of unique genes in each copy. This pattern suggests functional diversification among different ICEs, likely resulting from recombination and gene exchange processes occurring after insertion into the host (meter ref), reinforcing their role as dynamic reservoirs of adaptive functions. These genes were found to be associated with ParB/ThiF partitioning systems, which have been previously implicated in the partitioning of these types of elements in other bacteria (Acuña et al., 2013; Hickey et al., 2012) and toxin-antitoxin systems (TA). The system in question has been identified as belonging to the VapC/VapB category. These systems have been previously described in mobile elements, such as ICEclc from *Pseudomonas knackmussii*, where TA systems contribute to post-transfer maintenance and regulation of the element’s life cycle through a process called post-segregational killing (Miyazaki & van der Meer, 2011).

A notable proportion of the accessory genes exclusive to *S. granuli* TFA were implicated in metabolic adaptation and environmental resistance, characteristics that were conserved in the *Sphingobium baderi* species. The preservation of this accessory genome among species of different genera, despite their ecological divergence, lends support to the hypothesis of a recent and adaptive horizontal exchange of functional regions implicated in tolerance to toxic compounds and fine metabolic regulation. This phenomenon has been observed in other members of Sphingomonadales, where the transfer of ICEs has been shown to favor the expansion of aromatic metabolism and the response to environmental stress (Zhao et al., 2017). The presence of multiple genes encoding enzymes associated with the catabolism of aromatic compounds, such as dienelactone hydrolase, epoxide hydrolase, crotonase, and methylcrotonoyl–CoA carboxylase, suggests that these ICEs maintain the ability to degrade intermediates derived from aromatic hydrocarbons and polycyclic compounds, in line with the known degradative capacity of *S. granuli* TFA, as described in other Proteobacteria, such as Burkholderiales (Pérez-Pantoja et al., 2012). Furthermore, the detection of resistance genes, such as RND transporters, HlyD, and small multidrug resistance proteins, in conjunction with MarR-type regulators, suggests a coevolution of detoxification mechanisms and transcriptional control to optimize survival under chemical stress (Nishino & Yamaguchi, 2001).

It is important to emphasize the substantial number of variants present in the pangenome of integration sites, restriction-modification systems, and sigma-70 factors. This observation indicates that these elements have not only undergone a process of co-adaptation between the mobile element and its host (Oliveira et al., 2014), but can also modulate their activity in response to specific environmental conditions in a highly specific manner in each copy of each species. Furthermore, the presence of genes related to diverse metabolic pathways suggests that ICEs could contribute directly to the functional expansion of the host cell, facilitating its adaptation to variable ecological niches (Cury et al., 2017).

Finally, a comparative differential expression analysis was performed based on data from previous studies, allowing the basal condition to be compared with different types of metabolic, oxidative, and nitrosative stress. Despite the limitations in the design of the transcriptomic experiments that were analyzed, which were not specifically developed to study ICE3, the observed expression patterns provide a comprehensive framework for analyzing the transcriptomic plasticity contributed by both ICEs. The generalized induction of genes encoding universal stress proteins under most conditions acted as an internal positive control, thereby confirming the robustness and sensitivity of the experimental system for detecting global physiological responses. The functional differentiation previously observed between ICE6 and ICE3 is also reflected in the expression patterns observed by RNA-seq, where ICE3 genes showed greater activation under conditions of oxidative stress, nitrosative stress, or energy limitation, while those of ICE6 tended to remain transcriptionally stable.

In the absence of oxygen, a coordinated transcriptional pattern was observed involving the expression of genes related to maintaining redox balance. These genes included cytochrome bd oxidase, ferritin, thioredoxin peroxidase, and ArsR-type regulators. Despite the extensive research on the association of this gene set with adaptation to microaerophilic environments and protection against oxidative stress (Imlay, 2013), its metabolic significance in ICEs remains to be elucidated. By contrast, when the carbon source is altered from tetralin to β-hydroxybutyrate (BHB), a significant repression of these genes is observed, indicating a metabolic shift towards increased stability and reduced reliance on oxidative defense mechanisms. This transition is accompanied by a dynamic adjustment of respiratory machinery and antioxidant defense systems, suggesting a metabolic adaptation that enhances resilience in the face of oxidative stress. This change indicates that tetralin metabolism induces a compensatory redox response, possibly associated with the oxidation of aromatic compounds and the generation of reactive intermediates (González-Flores et al., 2019).

It has been observed that the AbiE toxin-antitoxin system present in ICE3 exhibited dynamic regulation that is contingent upon the stress condition. In the presence of nitrosative stress, the expression of the AbiEii toxin was observed, which has the potential to induce a state of partial persistence and to decelerate cell growth (Yang & Walsh, 2017). This finding, in synergy with the expression of genes encoding ABC transporters, enhances survival against reactive nitrogen species. These systems have been previously described in other Gram-negative bacteria as being pivotal to maintaining cell integrity under nitrosative stress conditions (Fetar et al., 2011). The AbiE system has also been implicated in the adaptation of the bacterium during nutrient limitation (stressfest) through the inhibition of the AbiEi antitoxin, thereby enabling toxin action and regulating cell growth. However, under nutrient conditions conducive to bacterial proliferation, AbiEi was induced, thereby enabling normal growth. These findings suggest that the AbiE system functions as a sensor and modulator of cell physiology, adjusting toxin activity according to the type and intensity of stress. This highlights its potential role in the resilience and adaptability of ICE3 and the host cell. This regulatory mechanism has been previously documented in other bacterial species (Chen et al., 2021).

In conclusion, this work provides a comprehensive overview of the mobile genetic elements of *Sphingopyxis granuli* TFA using complementary genomic and transcriptomic analysis approaches. Most of the identified ICEs appear to be incomplete, suggesting an evolutionary history marked by recombination and gene loss processes. However, ICE3 and ICE6 are noteworthy for their ability to conserve a repertoire of potentially functional cargo genes associated with detoxification, transport, and stress response mechanisms. These genes could confer adaptive advantages in variable environments. In particular, transcriptomic evidence indicates that ICE3 does not act as a passive element but could modulate the physiology of the host and contribute to its ability to resist multiple types of stress. This suggests that ICEs constitute active evolutionary platforms in bacterial adaptation that promote functional innovation and metabolic diversification in environmentally versatile bacteria such as *Sphingopyxis granuli* TFA.

## Acknowledgments

We would like to thank C3UPO and CICA for their HPC support. This work has been supported by the Grant PID2021-125491NB-I00 funded by MICIU/AEI/10.13039/501100011033 (Agencia Estatal de Investigación / Ministry of Science and Innovation of the Spanish Government), and by FEDER (UE) to F.R.R; and by the postdoctoral contract for I.G.R (PAIDI 2020, POSTDOC_21_00064) funded by Andalusian Government (Junta de Andalucía). A.R. and A.J.P. are funded by PID2023-150077OB-I00/AEI/10.13039/501100011033/ FEDER,UE (Agencia Estatal de Investigación / Ministry of Science and Innovation of the Spanish Government) and Plan Propio de Investigación y Transferencia (2023-2026) de la UPO, por la Consejería de Universidades, Investigación e Innovación de la Junta de Andalucía y por el Programa FEDER Andalucía 2021-2027, 2023/00002/014. A.M.R. is supported by doctoral fellowship PRE2022-104318, from the Agencia Estatal de Investigación, Ministerio de Ciencia e Investigación.

## References

Acuña, L. G., Cárdenas, J. P., Covarrubias, P. C., Haristoy, J. J., Flores, R., Nuñez, H., Riadi, G., Shmaryahu, A., Valdés, J., Dopson, M., Rawlings, D. E., Banfield, J. F., Holmes, D. S., & Quatrini, R. (2013). Architecture and gene repertoire of the flexible genome of the extreme acidophile Acidithiobacillus caldus. PloS One, 8(11), e78237. 10.1371/journal.pone.0078237

Anders, S., Pyl, P. T., & Huber, W. (2015). HTSeq—A Python framework to work with high-throughput sequencing data. Bioinformatics, 31(2), 166–169. 10.1093/bioinformatics/btu638

Asaf, S., Numan, M., Khan, A. L., & Al-Harrasi, A. (2020). Sphingomonas: From diversity and genomics to functional role in environmental remediation and plant growth. Critical Reviews in Biotechnology, 40(2), 138–152. 10.1080/07388551.2019.1709793

Bairoch, A., Apweiler, R., Wu, C. H., Barker, W. C., Boeckmann, B., Ferro, S., Gasteiger, E., Huang, H., Lopez, R., Magrane, M., Martin, M. J., Natale, D. A., O’Donovan, C., Redaschi, N., & Yeh, L.-S. L. (2005). The Universal Protein Resource (UniProt). Nucleic Acids Research, 33(Database Issue), D154-D159. 10.1093/nar/gki070

Bálint, B., Merényi, Z., Hegedüs, B., Grigoriev, I. V., Hou, Z., Földi, C., & Nagy, L. G. (2024). ContScout: Sensitive detection and removal of contamination from annotated genomes. Nature Communications, 15(1), 936. 10.1038/s41467-024-45024-5

Bean, E. L., Herman, C., Anderson, M. E., & Grossman, A. D. (2022). Biology and engineering of integrative and conjugative elements: Construction and analyses of hybrid ICEs reveal element functions that affect species-specific efficiencies. PLoS Genetics, 18(5), e1009998. 10.1371/journal.pgen.1009998

Bellanger, X., Payot, S., Leblond-Bourget, N., & Guédon, G. (2014). Conjugative and mobilizable genomic islands in bacteria: Evolution and diversity. FEMS Microbiology Reviews, 38(4), 720–760. 10.1111/1574-6976.12058

Bushnell, B. (2014). BBMap: A Fast, Accurate, Splice-Aware Aligner (LBNL-7065E). Ernest Orlando Lawrence Berkeley National Laboratory, Berkeley, CA (US). https://www.osti.gov/biblio/1241166

Camacho, C., Coulouris, G., Avagyan, V., Ma, N., Papadopoulos, J., Bealer, K., & Madden, T. L. (2009). BLAST+: Architecture and applications. BMC Bioinformatics, 10(1), 421. 10.1186/1471-2105-10-421

Cantalapiedra, C. P., Hernández-Plaza, A., Letunic, I., Bork, P., & Huerta-Cepas, J. (2021). eggNOG-mapper v2: Functional Annotation, Orthology Assignments, and Domain Prediction at the Metagenomic Scale. Molecular Biology and Evolution, 38(12), 5825–5829. 10.1093/molbev/msab293

Capra, J. A., Stolzer, M., Durand, D., & Pollard, K. S. (2013). How old is my gene? Trends in genetics : TIG, 29(11), 10.1016/j.tig.2013.07.001. 10.1016/j.tig.2013.07.001

Casimiro-Soriguer, C. S., Muñoz-Mérida, A., & Pérez-Pulido, A. J. (2017). Sma3s: A universal tool for easy functional annotation of proteomes and transcriptomes. Proteomics, 17(12). 10.1002/pmic.201700071

Che, Y., Wu, R., Li, H., Wang, L., Wu, X., Chen, Q., Chen, R., & Zhou, L. (2024). Molecular characterization of the integrative and conjugative elements harbouring multidrug resistance genes in Glaesserella parasuis. Veterinary Microbiology, 291, 110014. 10.1016/j.vetmic.2024.110014

Chen, X., Hu, A., Zou, Q., Luo, S., Wu, H., Yan, C., Liu, T., He, D., Li, X., & Cheng, G. (2021). The Mesorhizobium huakuii transcriptional regulator AbiEi plays a critical role in nodulation and is important for bacterial stress response. BMC Microbiology, 21, 245. 10.1186/s12866-021-02304-0

Christie, P. J., Whitaker, N., & González-Rivera, C. (2014). Mechanism and structure of the bacterial type IV secretion systems. Biochimica Et Biophysica Acta, 1843(8), 1578–1591. 10.1016/j.bbamcr.2013.12.019

Cury, J., Touchon, M., & Rocha, E. P. C. (2017). Integrative and conjugative elements and their hosts: Composition, distribution and organization. Nucleic Acids Research, 45(15), 8943–8956. 10.1093/nar/gkx607

de Assis, J. C. S., Gonçalves, O. S., Fernandes, A. S., de Queiroz, M. V., Bazzolli, D. M. S., & Santana, M. F. (2022). Genomic analysis reveals the role of integrative and conjugative elements in plant pathogenic bacteria. Mobile DNA, 13(1), 19. 10.1186/s13100-022-00275-1

de Dios, R., Rivas-Marin, E., Santero, E., & Reyes-Ramírez, F. (2020). Two paralogous EcfG σ factors hierarchically orchestrate the activation of the General Stress Response in Sphingopyxis granuli TFA. Scientific Reports, 10(1), 5177. 10.1038/s41598-020-62101-z

De Maayer, P., Chan, W.-Y., Martin, D. A. J., Blom, J., Venter, S. N., Duffy, B., Cowan, D. A., Smits, T. H. M., & Coutinho, T. A. (2015). Integrative conjugative elements of the ICEPan family play a potential role in Pantoea ananatis ecological diversification and antibiosis. Frontiers in Microbiology, 6, 576. 10.3389/fmicb.2015.00576

Delavat, F., Miyazaki, R., Carraro, N., Pradervand, N., & van der Meer, J. R. (2017). The hidden life of integrative and conjugative elements. FEMS Microbiology Reviews, 41(4), 512–537. 10.1093/femsre/fux008

Doron, S., Melamed, S., Ofir, G., Leavitt, A., Lopatina, A., Keren, M., Amitai, G., & Sorek, R. (2018). Systematic discovery of antiphage defense systems in the microbial pangenome. Science (New York, N.Y.), 359(6379), eaar4120. 10.1126/science.aar4120

Drobiazko, A., Adams, M. C., Skutel, M., Potekhina, K., Kotovskaya, O., Trofimova, A., Matlashov, M., Yatselenko, D., Maxwell, K. L., Blower, T. R., Severinov, K., Ghilarov, D., & Isaev, A. (2025). Molecular basis of foreign DNA recognition by BREX anti-phage immunity system. Nature Communications, 16(1), 1825. 10.1038/s41467-025-57006-2

Dy, R. L., Przybilski, R., Semeijn, K., Salmond, G. P. C., & Fineran, P. C. (2014). A widespread bacteriophage abortive infection system functions through a Type IV toxin-antitoxin mechanism. Nucleic Acids Research, 42(7), 4590–4605. 10.1093/nar/gkt1419

Fetar, H., Gilmour, C., Klinoski, R., Daigle, D. M., Dean, C. R., & Poole, K. (2011). mexEF-oprN multidrug efflux operon of Pseudomonas aeruginosa: Regulation by the MexT activator in response to nitrosative stress and chloramphenicol. Antimicrobial Agents and Chemotherapy, 55(2), 508–514. 10.1128/AAC.00830-10

García-Romero, I., Nogales, J., Díaz, E., Santero, E., & Floriano, B. (2020a). Understanding the metabolism of the tetralin degrader Sphingopyxis granuli strain TFA through genome-scale metabolic modelling. Scientific Reports, 10, 8651. 10.1038/s41598-020-65258-9

García-Romero, I., Nogales, J., Díaz, E., Santero, E., & Floriano, B. (2020b). Understanding the metabolism of the tetralin degrader Sphingopyxis granuli strain TFA through genome-scale metabolic modelling. Scientific Reports, 10, 8651. 10.1038/s41598-020-65258-9

García-Romero, I., Pérez-Pulido, A. J., González-Flores, Y. E., Reyes-Ramírez, F., Santero, E., & Floriano, B. (2016). Genomic analysis of the nitrate-respiring Sphingopyxis granuli (formerly Sphingomonas macrogoltabida) strain TFA. BMC Genomics, 17, 93. 10.1186/s12864-016-2411-1

Garcillán-Barcia, M. P., Cuartas-Lanza, R., Cuevas, A., & de la Cruz, F. (2019). Cis-Acting Relaxases Guarantee Independent Mobilization of MOBQ 4 Plasmids. Frontiers in Microbiology, 10, 2557. 10.3389/fmicb.2019.02557

Garcillán-Barcia, M. P., Francia, M. V., & de La Cruz, F. (2009). The diversity of conjugative relaxases and its application in plasmid classification. FEMS Microbiology Reviews, 33(3), 657–687. 10.1111/j.1574-6976.2009.00168.x

Ghinet, M. G., Bordeleau, E., Beaudin, J., Brzezinski, R., Roy, S., & Burrus, V. (2011). Uncovering the prevalence and diversity of integrating conjugative elements in actinobacteria. PloS One, 6(11), e27846. 10.1371/journal.pone.0027846

Gomberg, A. F., & Grossman, A. D. (2024). It’s complicated: Relationships between integrative and conjugative elements and their bacterial hosts. Current Opinion in Microbiology, 82, 102556. 10.1016/j.mib.2024.102556

González-Flores, Y. E., de Dios, R., Reyes-Ramírez, F., & Santero, E. (2019). The response of Sphingopyxis granuli strain TFA to the hostile anoxic condition. Scientific Reports, 9, 6297. 10.1038/s41598-019-42768-9

Guédon, G., Charron-Bourgoin, F., Lacroix, T., Hamadouche, T., Soler, N., Douzi, B., Chiapello, H., & Leblond-Bourget, N. (2025). Massive acquisition of conjugative and mobilizable integrated elements fuels Faecalibacterium plasticity and hints at their adaptation to the gut. Scientific Reports, 15(1), 17013. 10.1038/s41598-025-99981-y

Guédon, G., Libante, V., Coluzzi, C., Payot, S., & Leblond-Bourget, N. (2017). The Obscure World of Integrative and Mobilizable Elements, Highly Widespread Elements that Pirate Bacterial Conjugative Systems. Genes, 8(11), 337. 10.3390/genes8110337

Haskett, T. L., Ramsay, J. P., Bekuma, A. A., Sullivan, J. T., O’Hara, G. W., & Terpolilli, J. J. (2017). Evolutionary persistence of tripartite integrative and conjugative elements. Plasmid, 92, 30–36. 10.1016/j.plasmid.2017.06.001

He, J., Yang, Z., Wang, M., Jia, R., Chen, S., Liu, M., Zhao, X., Yang, Q., Wu, Y., Zhang, S., Huang, J., Ou, X., Sun, D., Tian, B., He, Y., Wu, Z., Cheng, A., & Zhu, D. (2024). Integrative and conjugative elements of Pasteurella multocida: Prevalence and signatures in population evolution. Virulence, 15(1), 2359467. 10.1080/21505594.2024.2359467

Hernández-Arriaga, A. M., Rubio-Lepe, T. S., Espinosa, M., & del Solar, G. (2009). Repressor CopG prevents access of RNA polymerase to promoter and actively dissociates open complexes. Nucleic Acids Research, 37(14), 4799–4811. 10.1093/nar/gkp503

Hickey, W. J., Chen, S., & Zhao, J. (2012). The phn Island: A New Genomic Island Encoding Catabolism of Polynuclear Aromatic Hydrocarbons. Frontiers in Microbiology, 3, 125. 10.3389/fmicb.2012.00125

Hollingshead, S., McVicker, G., Nielsen, M. R., Zhang, Y., Pilla, G., Jones, R. A., Thomas, J. C., Johansen, S. E. H., Exley, R. M., Brodersen, D. E., & Tang, C. M. (2024). Shared mechanisms of enhanced plasmid maintenance and antibiotic tolerance mediated by the VapBC toxin:antitoxin system. mBio, 16(2), e02616–24. 10.1128/mbio.02616-24

Imlay, J. A. (2013). The molecular mechanisms and physiological consequences of oxidative stress: Lessons from a model bacterium. Nature Reviews Microbiology, 11(7), 443–454. 10.1038/nrmicro3032

Johnson, C. M., & Grossman, A. D. (2015). Integrative and Conjugative Elements (ICEs): What They Do and How They Work. Annual Review of Genetics, 49, 577–601. 10.1146/annurev-genet-112414-055018

Johnson, C. M., Harden, M. M., & Grossman, A. D. (2022). Interactions between mobile genetic elements: An anti-phage gene in an integrative and conjugative element protects host cells from predation by a temperate bacteriophage. PLOS Genetics, 18(2), e1010065. 10.1371/journal.pgen.1010065

Jones, P., Binns, D., Chang, H.-Y., Fraser, M., Li, W., McAnulla, C., McWilliam, H., Maslen, J., Mitchell, A., Nuka, G., Pesseat, S., Quinn, A. F., Sangrador-Vegas, A., Scheremetjew, M., Yong, S.-Y., Lopez, R., & Hunter, S. (2014). InterProScan 5: Genome-scale protein function classification. Bioinformatics (Oxford, England), 30(9), 1236–1240. 10.1093/bioinformatics/btu031

Kalyaanamoorthy, S., Minh, B. Q., Wong, T. K. F., von Haeseler, A., & Jermiin, L. S. (2017). ModelFinder: Fast model selection for accurate phylogenetic estimates. Nature Methods, 14(6), Article 6. 10.1038/nmeth.4285

Katoh, K., & Standley, D. M. (2013). MAFFT multiple sequence alignment software version 7: Improvements in performance and usability. Molecular Biology and Evolution, 30(4), 772–780. 10.1093/molbev/mst010

Kim, D., Paggi, J. M., Park, C., Bennett, C., & Salzberg, S. L. (2019). Graph-based genome alignment and genotyping with HISAT2 and HISAT-genotype. Nature Biotechnology, 37(8), 907–915. 10.1038/s41587-019-0201-4

Long, X., Wang, X., Mao, D., Wu, W., & Luo, Y. (2022). A Novel XRE-Type Regulator Mediates Phage Lytic Development and Multiple Host Metabolic Processes in Pseudomonas aeruginosa. Microbiology Spectrum, 10(6), e0351122. 10.1128/spectrum.03511-22

Love, M. I., Huber, W., & Anders, S. (2014). Moderated estimation of fold change and dispersion for RNA-seq data with DESeq2. Genome Biology, 15(12), 550. 10.1186/s13059-014-0550-8

Luo, D., Wu, Z., Bai, Q., Zhang, Y., Huang, M., Huang, Y., & Li, X. (2023). Universal Stress Proteins: From Gene to Function. International Journal of Molecular Sciences, 24(5), 4725. 10.3390/ijms24054725

Makarova, K. S., Krupovic, M., & Koonin, E. V. (2014). Evolution of replicative DNA polymerases in archaea and their contributions to the eukaryotic replication machinery. Frontiers in Microbiology, 5. 10.3389/fmicb.2014.00354

Meygret, A., Peuchant, O., Dordet-Frisoni, E., Sirand-Pugnet, P., Citti, C., Bébéar, C., Béven, L., & Pereyre, S. (2019). High Prevalence of Integrative and Conjugative Elements Encoding Transcription Activator-Like Effector Repeats in Mycoplasma hominis. Frontiers in Microbiology, 10, 2385. 10.3389/fmicb.2019.02385

Millman, A., Melamed, S., Leavitt, A., Doron, S., Bernheim, A., Hör, J., Garb, J., Bechon, N., Brandis, A., Lopatina, A., Ofir, G., Hochhauser, D., Stokar-Avihail, A., Tal, N., Sharir, S., Voichek, M., Erez, Z., Ferrer, J. L. M., Dar, D., … Sorek, R. (2022). An expanded arsenal of immune systems that protect bacteria from phages. Cell Host & Microbe, 30(11), 1556–1569.e5. 10.1016/j.chom.2022.09.017

Minh, B. Q., Schmidt, H. A., Chernomor, O., Schrempf, D., Woodhams, M. D., von Haeseler, A., & Lanfear, R. (2020). IQ-TREE 2: New Models and Efficient Methods for Phylogenetic Inference in the Genomic Era. Molecular Biology and Evolution, 37(5), 1530–1534. 10.1093/molbev/msaa015

Miyazaki, R., & van der Meer, J. R. (2011). A dual functional origin of transfer in the ICEclc genomic island of Pseudomonas knackmussii B13. Molecular Microbiology, 79(3), 743–758. 10.1111/j.1365-2958.2010.07484.x

Nishino, K., & Yamaguchi, A. (2001). Analysis of a complete library of putative drug transporter genes in Escherichia coli. Journal of Bacteriology, 183(20), 5803–5812. 10.1128/JB.183.20.5803-5812.2001

Oliveira, P. H., Touchon, M., & Rocha, E. P. C. (2014). The interplay of restriction-modification systems with mobile genetic elements and their prokaryotic hosts. Nucleic Acids Research, 42(16), 10618–10631. 10.1093/nar/gku734

Pérez-Pantoja, D., Donoso, R., Agulló, L., Córdova, M., Seeger, M., Pieper, D. H., & González, B. (2012). Genomic analysis of the potential for aromatic compounds biodegradation in Burkholderiales. Environmental Microbiology, 14(5), 1091–1117. 10.1111/j.1462-2920.2011.02613.x

Russo, T. A., & Marr, C. M. (2019). Hypervirulent Klebsiella pneumoniae. Clinical Microbiology Reviews, 32(3), e00001–19. 10.1128/CMR.00001-19

Schwengers, O., Jelonek, L., Dieckmann, M. A., Beyvers, S., Blom, J., & Goesmann, A. (2021). Bakta: Rapid and standardized annotation of bacterial genomes via alignment-free sequence identification. Microbial Genomics, 7(11), 000685. 10.1099/mgen.0.000685

Seemann, T. (2013). Barrnap: BAsic Rapid Ribosomal RNA Predictor.

Sentchilo, V., Czechowska, K., Pradervand, N., Minoia, M., Miyazaki, R., & van der Meer, J. R. (2009). Intracellular excision and reintegration dynamics of the ICEclc genomic island of Pseudomonas knackmussii sp. Strain B13. Molecular Microbiology, 72(5), 1293–1306. 10.1111/j.1365-2958.2009.06726.x

Sharma, M., Khurana, H., Singh, D. N., & Negi, R. K. (2021). The genus Sphingopyxis: Systematics, ecology, and bioremediation potential - A review. Journal of Environmental Management, 280, 111744. 10.1016/j.jenvman.2020.111744

Shaw, J., & Yu, Y. W. (2023). Fast and robust metagenomic sequence comparison through sparse chaining with skani. Nature Methods, 20(11), 1661–1665. 10.1038/s41592-023-02018-3

Song, D., Chen, X., Yao, H., Kong, G., Xu, M., Guo, J., & Sun, G. (2023). The variations of native plasmids greatly affect the cell surface hydrophobicity of sphingomonads. mSystems, 8(6), e0086223. 10.1128/msystems.00862-23

Subramoni, S., & Venturi, V. (2009). LuxR-family «solos»: Bachelor sensors/regulators of signalling molecules. Microbiology (Reading, England), 155(Pt 5), 1377–1385. 10.1099/mic.0.026849-0

Sullivan, J. T., & Ronson, C. W. (1998). Evolution of rhizobia by acquisition of a 500-kb symbiosis island that integrates into a phe-tRNA gene. Proceedings of the National Academy of Sciences of the United States of America, 95(9), 5145–5149. 10.1073/pnas.95.9.5145

Tettelin, H., Masignani, V., Cieslewicz, M. J., Donati, C., Medini, D., Ward, N. L., Angiuoli, S. V., Crabtree, J., Jones, A. L., Durkin, A. S., DeBoy, R. T., Davidsen, T. M., Mora, M., Scarselli, M., Margarit y Ros, I., Peterson, J. D., Hauser, C. R., Sundaram, J. P., Nelson, W. C., … Fraser, C. M. (2005). Genome analysis of multiple pathogenic isolates of Streptococcus agalactiae: Implications for the microbial “pan-genome”. Proceedings of the National Academy of Sciences, 102(39), 13950–13955. 10.1073/pnas.0506758102

Tokuda, M., & Shintani, M. (2024). Microbial evolution through horizontal gene transfer by mobile genetic elements. Microbial Biotechnology, 17(1), e14408. 10.1111/1751-7915.14408

Tonkin-Hill, G., MacAlasdair, N., Ruis, C., Weimann, A., Horesh, G., Lees, J. A., Gladstone, R. A., Lo, S., Beaudoin, C., Floto, R. A., Frost, S. D. W., Corander, J., Bentley, S. D., & Parkhill, J. (2020). Producing polished prokaryotic pangenomes with the Panaroo pipeline. Genome Biology, 21(1), 180. 10.1186/s13059-020-02090-4

Touchon, M., Bobay, L.-M., & Rocha, E. P. (2014). The chromosomal accommodation and domestication of mobile genetic elements. Current Opinion in Microbiology, 22, 22–29. 10.1016/j.mib.2014.09.010

van Vliet, A. H. M., Charity, O. J., & Reuter, M. (2021). A Campylobacter integrative and conjugative element with a CRISPR-Cas9 system targeting competing plasmids: A history of plasmid warfare? Microbial Genomics, 7(11), 000729. 10.1099/mgen.0.000729

Wang, M., Liu, G., Liu, M., Tai, C., Deng, Z., Song, J., & Ou, H.-Y. (2024). ICEberg 3.0: Functional categorization and analysis of the integrative and conjugative elements in bacteria. Nucleic Acids Research, 52(D1), D732–D737. 10.1093/nar/gkad935

Wozniak, R. A. F., Fouts, D. E., Spagnoletti, M., Colombo, M. M., Ceccarelli, D., Garriss, G., Déry, C., Burrus, V., & Waldor, M. K. (2009). Comparative ICE genomics: Insights into the evolution of the SXT/R391 family of ICEs. PLoS Genetics, 5(12), e1000786. 10.1371/journal.pgen.1000786

Wozniak, R. A. F., & Waldor, M. K. (2010a). Integrative and conjugative elements: Mosaic mobile genetic elements enabling dynamic lateral gene flow. Nature Reviews. Microbiology, 8(8), 552–563. 10.1038/nrmicro2382

Wozniak, R. A. F., & Waldor, M. K. (2010b). Integrative and conjugative elements: Mosaic mobile genetic elements enabling dynamic lateral gene flow. Nature Reviews Microbiology, 8(8), 552–563. 10.1038/nrmicro2382

Yang, Q. E., & Walsh, T. R. (2017). Toxin–antitoxin systems and their role in disseminating and maintaining antimicrobial resistance. FEMS Microbiology Reviews, 41(3), 343–353. 10.1093/femsre/fux006

Yu, G., Smith, D. K., Zhu, H., Guan, Y., & Lam, T. T.-Y. (2017). ggtree: An r package for visualization and annotation of phylogenetic trees with their covariates and other associated data. Methods in Ecology and Evolution, 8(1), 28–36. 10.1111/2041-210X.12628

Zhao, Q., Yue, S., Bilal, M., Hu, H., Wang, W., & Zhang, X. (2017). Comparative genomic analysis of 26 Sphingomonas and Sphingobium strains: Dissemination of bioremediation capabilities, biodegradation potential and horizontal gene transfer. The Science of the Total Environment, 609, 1238–1247. 10.1016/j.scitotenv.2017.07.249

